# Chronic dietary ingestion of *Bacillus thuringiensis* spores promotes intestinal inflammation and aging in non-target adult *Drosophila*

**DOI:** 10.1101/2025.03.25.645302

**Authors:** Aurélia Joly, Julie Soltys, Jade Finkelstein, Corinne Rancurel, Audrey Amate, Marie-Paule Nawrot-Esposito, Benoit Chassaing, Armel Gallet, Raphaël Rousset

## Abstract

*Bacillus thuringiensis* (*Bt*), a spore-forming Gram-positive bacterium, is the leading microbial insecticide used in both organic and conventional agriculture to fight lepidopteran larvae. *Bt* insecticides, which consist of a mix of *Bt* spores and crystals of entomopathogenic toxins (Cry toxins), kill target pest larvae rapidly after ingestion by destroying their intestinal epithelium. The potential adverse effects of chronic consumption of *Bt* spores by non-target organisms have not been thoroughly investigated. In this study, using adult *Drosophila melanogaster*, a non-target dipteran organism of *Bt* insecticides, we show that chronic ingestion of agricultural doses of spores in the diet reduces lifespan. Our results demonstrate that *Bt* spore ingestion affects gut morphology, promotes dysplasia, alters septate junctions and enhances epithelial permeability. In addition, we observed increased levels of inflammatory signaling pathways and reactive oxygen species. Taken together, our results indicate that chronic consumption of *Bt* spores promotes inflammation and oxidative stress, leading to premature aging of the gut and early lethality in *Drosophila*. Extended to non-target insects, which account for 85% of animal biodiversity, our study suggests that highly persistent *Bt* spores may have unintended long-term effects on the environment.

## INTRODUCTION

The use of pesticides exploded in the mid-20th century to meet the world’s growing food needs and the demand for profitable crops. The massive use of these chemical substances over several decades has had a dramatic impact on ecology, and the effects of some of them on diseases have also been reported (1). To reduce the use of chemical pesticides, alternative methods of plant protection using natural mechanisms, known as biocontrol, have emerged. Among these solutions, the spores of the bacterium *Bacillus thuringiensis* (*Bt*) are widely used as an insecticide against pest larvae that ravage crops (tomatoes, cabbage, lettuce, corn, etc.) and forests (pine trees), but also against harmful insects such as mosquitoes (2). *Bt* accounts for 32000 tons of products sold in both organic and conventional agriculture, making it the second most widely used insecticide (chemical and microbial combined) (3). Moreover, the use of *Bt* insecticides is expected to continue to grow due to government and societal incentives to promote biocontrol solutions as safer, more sustainable alternatives to chemical pesticides.

*Bt* belongs to the *Bacillus cereus* (*Bc*) group, a distinctive cluster of closely genetically related species within the genus *Bacillus* that are Gram-positive, facultative anaerobic, and ubiquitous in the environment (4). Under unfavorable conditions (low pH, nutrient scarcity), vegetative bacteria of this group have the ability to transform into spores, a resistant dormant state that ensures survival until conditions improve. Within the *Bc* group, *Bc sensu stricto* is an opportunistic food poisoning pathogen that causes diarrhea involving the enterotoxins Nhe, Hbl and CytK. *Bc sensu stricto* infection is the leading cause of foodborne outbreaks (FBO) among toxin-producing bacteria, and second behind *Salmonella* in total causative agents in Europe (5). Some studies have further highlighted the lethal potential of *Bc* in immunosuppressed individuals (6, 7). In addition to the insecticidal properties of *Bt*, *Bc* group bacteria exhibit other attributes that can be exploited in biocontrol (antibiotic, antifungal and nematicidal activities), plant growth promotion (biofertilizers) and human and veterinary medicine (probiotics) (8-11). This highlights the complexity of the *Bc* group, which displays a variety of beneficial and pathogenic properties.

The insecticidal properties of *Bt* are conferred by the enthomopathogenic crystal toxins (Cry toxins) produced during sporulation. Each *Bt* subspecies produces a unique Cry toxin cocktail that confers specific insecticidal properties able to target an unique insect order, a characteristic used in biocontrol solutions (12, 13). After dispersion, *Bt* insecticides (a mix of spores and toxin crystals) are ingested by insects at the larval stage and enter the digestive tract, which provides favorable conditions for spore germination and toxin release. Once dissolved and activated by intestinal proteases, the toxins bind to intestinal receptors such as Cadherins, Alkaline phosphatases, Aminopeptidases and/or ABCC transporters present on the surface of the epithelium and create pores in the intestinal cells (13-15). The bacteria present in the lumen can invade the systemic circulation and cause death of the larvae by septicemia (16). Importantly, no acute toxicity of *Bt* products has been reported in mammals, but the long-term effects of ingesting low doses are poorly documented (17-23). Given the prominent and increasing use of these persistent spore-based insecticides, there is an urgent need to fill the knowledge gap and investigate their potential adverse effects on non-target organisms when chronically ingested with food (23). Three recent studies have shown that several strains of *Bt* from agricultural products have been traced back to foodborne outbreaks, suggesting that *Bt* may pose a risk to food safety (24-26).

Previous work in the non-target organism *Drosophila* reported that low doses of *Bt* vegetative bacteria induced a transient hyperplasia dependent on the JNK, JAK/STAT and Hippo/Yki signaling pathways in adults (27). With regard to the ingestion of *Bt* spores, it has been shown in fruit fly larvae that growth defects and developmental delays occur, mainly due to the death of enterocytes, which impairs protein digestion, and mortality at higher doses. (28, 29). In rats, *Bt* spores can germinate and persist in the intestine for several days (30), and it has recently been reported that persistence of *Bt* spores occurs in the posterior part of the adult *Drosophila* midgut due to a negative feedback mechanism that dampens the immune response (31). In mammals, microbiome dysbiosis and growth of pathogenic bacteria in the intestine are known to induce a pro-inflammatory state that can lead to the development of Intestinal Bowel Diseases (IBD) such as Crohn’s disease or ulcerative colitis and, in the worst cases, cancer (32, 33). Yet, the potential consequences of chronic dietary intake of *Bt* spores on the development of intestinal pathophysiology are not known. Here, we used the adult D*rosophila melanogaster* intestine as a model to assess the effects of chronic ingestion of low doses of *Bt* spores on non-target insect species.

## RESULTS

### Chronic ingestion of low-dose *Bt* spores with the diet reduces *Drosophila* lifespan

We exposed mated *Drosophila melanogaster* females, a non-target organism for the insecticidal properties of *Bt*, to commercial formulations containing *Bt* subspecies *kurstaki* (*Btk,* used worldwide as a microbial insecticide to kill lepidopteran larvae) along with their regular diet. To this end, we developed a standardized protocol in which tubes containing food and *Bt* product were changed every 3-4 days prior to spore germination on the medium (31), thus reproducing regular dietary exposure to *Bt* spores only. The recommendations for the spraying of *Bt* products in France and Europe are 1 to 1.5 kg/ha (corresponding to ~4 to 6x10^5^ colony forming units (CFU)/tube, see Materials and Methods) per application, with 2 to 12 repetitions ([EU Pesticides Database](https://food.ec.europa.eu/plants/pesticides/eu-pesticides-database_en)). We initially exposed flies to a dose of 10^6^ CFU/tube/fly, corresponding to approximately two consecutive treatments. The commercial products DiPel and Delfin contain the closely related *Btk* strains ABTS-351 and SA-11, respectively. We observed that exposure to both commercial products at the dose of 10^6^ significantly reduced the lifespan of the flies (Fig. 1A; Fig. S1A). To confirm that the flies were only ingesting spores and not vegetative bacteria that might have germinated on the medium, we reiterated the experiment and changed the tubes on a daily basis. We obtained a similar significant reduction in fly lifespan, confirming that this effect was not due to vegetative forms of the bacteria (Fig. S1B). In addition to spores and crystals, commercial formulations contain additives to improve dispersion, adhesion and preservation of the product. To test the possibility that the lifespan is affected by additives, we produced laboratory-made spores of *Btk* ABTS-351 or SA-11 without adjuvants. We observed that the chronic ingestion of these additive-free spores had the same effect as the full formulation of DiPel and Delfin products. These results assess that *Bt* spores, and not *Bt* vegetative bacteria or additives, are solely responsible for the increased mortality observed in flies (Fig. 1B; Fig. S1C). Moreover, we tested whether the Cry toxins produced by the spores were responsible for this deleterious effect. We used the acrystalliferous derivative of the SA-11 strain (SA-11ΔCry) (31), which does not produce Cry toxins, and still observed early lethality, supporting the deleterious effect of the spores alone (Fig. 1F).

**Figure 1.**
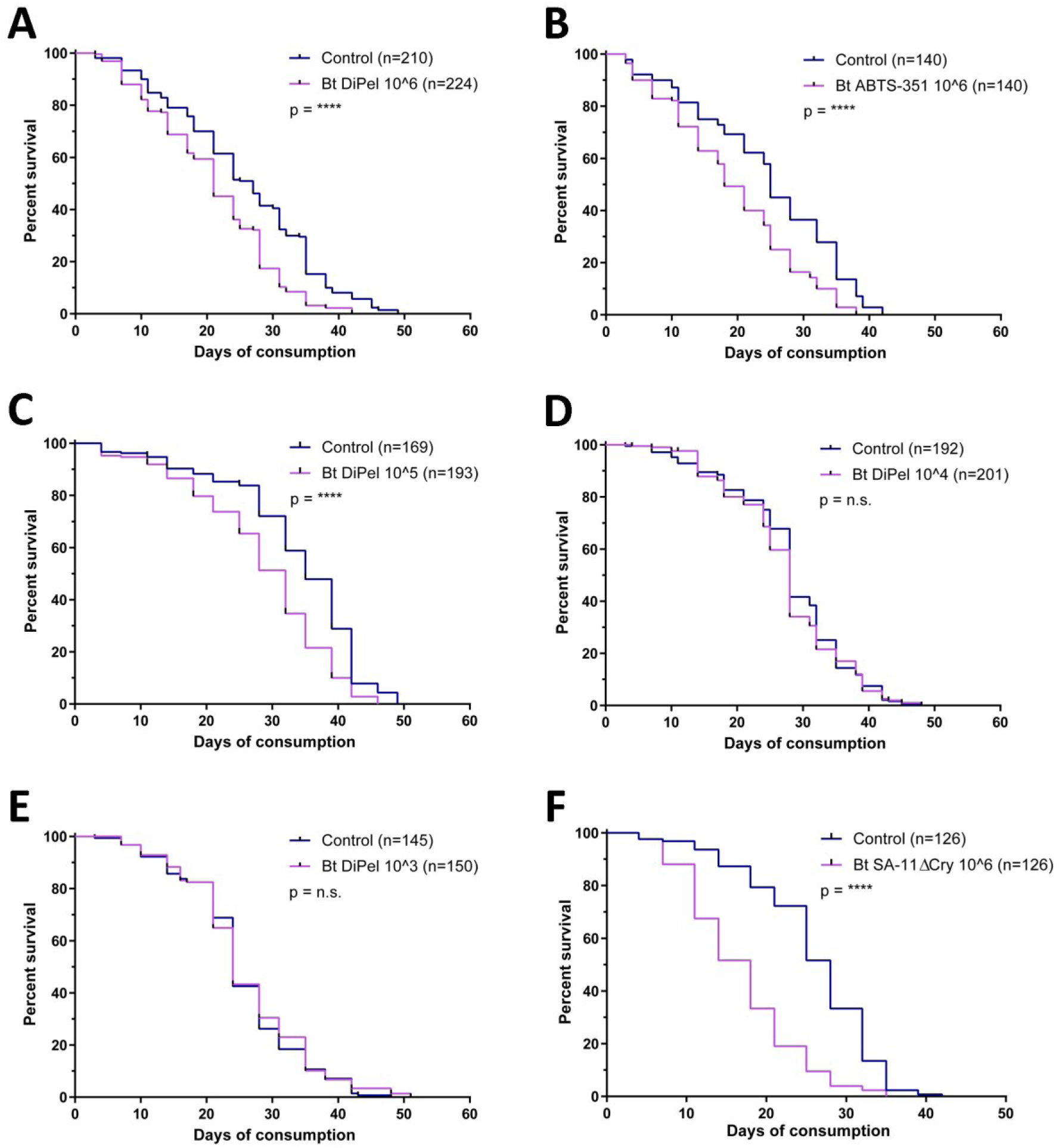
*Drosophila* survival during chronic ingestion of low doses of *Bt* spores with the diet. **A**) Survival assay at a dose of 10^6^ CFU/*Drosophila*/tube of DiPel starting on day 0 of treatment. **B)** Survival assay with homemade *Bt* spores ABTS-351 (same strain as the DiPel product) at dose 10^6^. **C**-**E)** Survival assays at the indicated concentrations (10^5^, 10^4^, 10^3^). **F)** Survival assay with homemade *Bt* spores SA-11 ΔCry (acrystalliferous form) at dose 10^6^. *n is indicated in each panel. Results are presented with Kaplan-Meier curves and statistics are Log-Rank Mantel-Cox tests*.

We next investigated the effect of lower doses to assess the impact on lifespan in a dose-dependent manner. To do this, we performed survival assays with doses of DiPel ranging from 10^5^ to 10^3^ CFU/tube/fly (Fig. 1C-E). We observed that a dose of 10^5^ CFU/tube/fly, 10 times lower than previously used, was sufficient to significantly reduce the fly lifespan. However, the lower doses of 10^4^ and 10^3^ CFU/tube/fly did not significantly affect the lifespan of *Bt*-exposed flies. Taken together, these results demonstrate that chronic ingestion of low doses of *Bt* spores has a critical impact on the lifespan of the non-target organism *Drosophila*, independently of the commercial additives and Cry toxins.

### Spore germination is not sufficient for early lethality

Numerous studies have already shown that after ingestion of vegetative bacteria, the innate immune response leads to their rapid and efficient elimination in the anterior part of the intestine (34). In contrast, we have recently shown that spores ingested by *Drosophila* pass through the anterior part of the intestine without eliciting an immune response (31). The spores then reach the posterior part of the intestine where they find a suitable environment for germination, leading to persistence of *Bt* up to 10 days after ingestion. Therefore, we wanted to address whether the presence of vegetative bacteria, due to spore germination in the posterior part of the intestine, is sufficient to cause the early lethality we observed previously. We first analyzed the germination profile (i.e. bacterial load of vegetative forms versus spores) of different *Bacillus* strains *in vivo* in the *Drosophila* gut over time during chronic ingestion. We performed CFU quantification from whole midguts by comparing heat-treated intestinal extracts (to kill germinating spores and vegetative cells, but not the spores) to non-heat-treated samples (total, i.e. spores, germinating spores and vegetative cells). In this experiment, we analyzed the germination profile of *Bt* SA-11, the toxin-free *Bt* SA-11ΔCry strain, and *Bc* (reference strain ATCC 14579). In addition, we used the strain *Bacillus amyloliquefaciens,* which is used as a biofertilizer in agriculture and is considered to have no harmful side effects (35). The results show for all strains that *Bacillus* spores germinate in the gut, that both spore and vegetative forms are present at comparable levels, and that the bacterial load remains relatively constant over time (Fig. 2A-D). To test if the presence of the vegetative forms of *Bacillus* causes the early lethality observed previously, we performed a survival assay comparing the survival probability between the pathogenic strain *Bc* ATCC 14579 and the harmless strain *Bacillus amyloliquefaciens.* While we observed a reduction in longevity with the pathogenic *Bc* strain comparable to what we previously observed with *Bt* (Fig. 2E), we did not observe any effect on *Drosophila* lifespan with *Bacillus amyloliquefacien* (Fig. 2F). Altogether, these results indicate that spore germination in the gut is not sufficient to cause increased lethality, but that the virulence of the strain is a critical factor (36).

**Figure 2.**
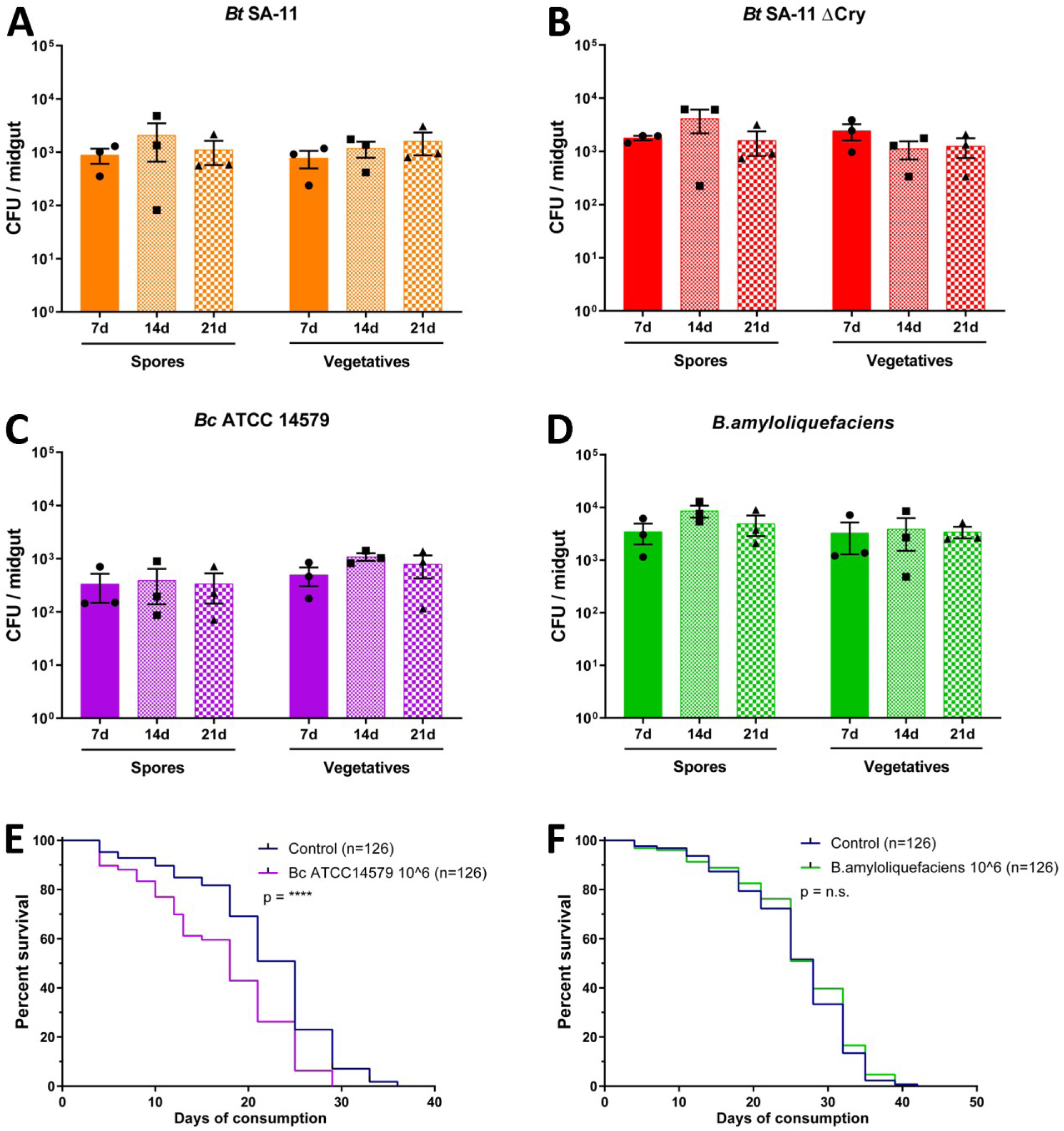
Bacterial load and *Drosophila* survival upon chronic ingestion of *Bacillus* spores A-D) Number of spores and vegetative cells per gut at three time points 7d, 14d, 21d after chronic ingestion of *Bt* SA-11 (A), *Bt* SA-11ΔCry (B), *Bc* ATCC14579 (C) and *Bacillus amyloliquefaciens* (D) spores at dose 10^6^. *Three independent replicates were performed with a total of 15 guts (n) for each strain*. **E-F)** Survival after chronic ingestion of *Bc* ATCC14579 (E) and *Bacillus amyloliquefaciens* (F) spores. *Kaplan-Meier curves are presented (n is indicated in each panel) and statistics are Log-Rank Mantel-Cox tests*.

### Transcriptome analysis reveals inflammation and oxidative stress in *Bt*-challenged flies

We performed a transcriptomic analysis of fly midguts after 7, 14 and 21 days of chronic *Bt* consumption to investigate the intestinal epithelial response. In a chronic context, we expected that small changes in gene expression over time could be deleterious. For this reason, we considered in our analysis that an arbitrary threshold at 30% in gene expression variation among significant differentially expressed genes could significantly influence intestinal physiology in the long term. The RNAseq analysis shows a variation in gene expression that progresses over time with chronic exposure to a low dose of *Bt* spores (Fig. 3A, B). A total of 342 differentially expressed genes (adjusted p-value < 0.05) with a fold change of at least 1.3 or less than -1.3 (i.e. threshold of 30%) in the three time points were identified. Among these genes, 193 are upregulated and 151 are downregulated (Fig. 3B). The differentially expressed genes belong to several enriched pathways related to detoxification and oxidoreduction, especially glutathione and cytochrome p450, as well as general metabolic pathways, energy metabolism (sulfur), amino acid metabolism (alanine, aspartate, glutamate) and vitamin metabolism (folate biosynthesis) (Fig. 3C; Fig. S2A and B). Importantly, 9 genes related to glutathione metabolic process are strongly upregulated in the *Bt* conditions (*GstD5, GstD4, GstD6, GstD2, GstS1, GstD1, GstD9, GstE6, GstE7*) (Fig. 3D). These genes encode for glutathione S-transferase enzymes, which are part of the detoxification system in conjunction with cytochrome p450 proteins and of the antioxidant response elements (AREs). As a protective mechanism against oxidative stress, these AREs include other enzymes such as catalases, peroxiredoxins, thioredoxins, glutathione peroxidases and reductases, but also protein chaperones and components of the proteasome complex to repair or eliminate damaged proteins (37, 38). Interestingly, we found strong upregulation of ARE enzymes such as Thioredoxin reductase 1 (Trxr1), Thioredoxin 2 (Trx2), Peroxiredoxin 2 (Prx2), and Clot (Cl) (Fig. 3D). We identified other components known to be involved in the oxidative stress response and/or the oxidation-reduction process such as Naam (39), Ntc (40), Sni (41), Mbf1 (42), Atg18b (43) and AOX1 (44). In addition, the transcriptomic analysis revealed expression changes in several genes involved in lipid and carbohydrate metabolism such as the triglyceride lipase Bmm (45), the α-glucosidase Tobi (46) and the lactate dehydrogenase Ldh (47) (Fig. 3D), suggesting that the metabolism of *Bt*-treated flies may be modified.

**Figure 3.**
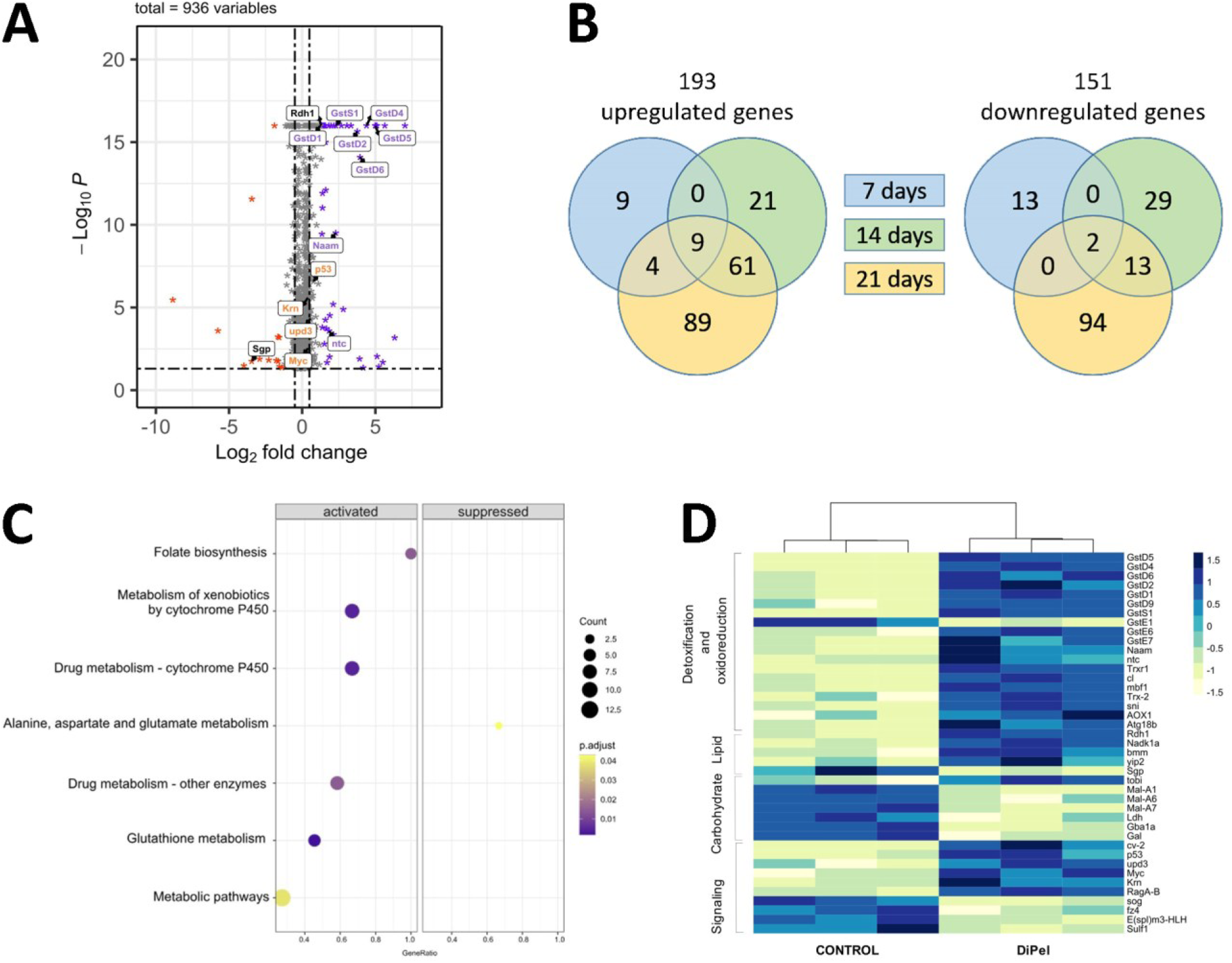
Transcriptomic analysis of intestines exposed to chronic ingestion of *Bt* spores. **A)** Volcano plot of differentially expressed genes in the *Drosophila* gut between *Bt* DiPel at 10^6^ CFU/*Drosophila*/tube and control (water). Each star represents a gene whose expression is significantly modified with a fold-change of less than -1.3 (orange) and more than +1.3 (purple) at all time points regrouped (7, 14 and 21 days of chronic ingestion). A set of genes of interest is indicated (see main text for details). **B)** Number of the differentially expressed genes shared between the three time points (Venn diagram). **C)** GO terms (KEGG pathways) found in the differentially expressed genes (three time points regrouped) showing enrichment of genes involved in detoxification/oxidation, carbohydrate/lipid metabolism and signaling. **D)** Heatmap showing normalized expression of key genes differentially expressed in these processes.

We also noticed that genes associated with important signaling pathways involved in intestinal inflammation, proliferation and regeneration were affected. We identified two upregulated ligands (Fig. 3D), Keren (Krn) and Unpaired3 (Upd3), of the EGFR and JAK-STAT pathways, respectively (48-52). We also observed an upregulation of the transcription factor Myc, which has been shown to act as an integrator of multiple signaling pathways to control intestinal stem cell proliferation during midgut regeneration (53). Moreover, data revealed an upregulation of p53 (Fig. 3D), which is required for adaptive responses to genotoxic stress, including cell death, compensatory proliferation and DNA repair mechanisms (54). Overall, the transcriptomic data indicate that a chronic *Bt* exposure results in a global gut response characterized by detoxification, oxidative stress, metabolic adaptation, inflammation and regeneration.

### Chronic *Bt* ingestion affects enterocyte morphology, alters septate junctions and increases gut permeability

We next explored how these transcriptomic changes affect the gut epithelium at the cellular level. To observe the general morphology of the epithelium after 14 days of chronic *Bt* consumption, we performed histological cross-sections of *Myo1A-GAL4,UAS-GFP* flies (expressing GFP in enterocytes). We observed that the regular pseudo-stratification of nuclei seen in control guts was disorganized in those challenged with *Bt* (Fig. 4A). Strikingly, we also noticed that the morphology of the enterocytes was altered, forming a distorted surface on contact with the lumen instead of a linear and regular one. These observations indicate that chronic *Bt* consumption affects the morphology of the intestinal epithelium.

**Figure 4.**
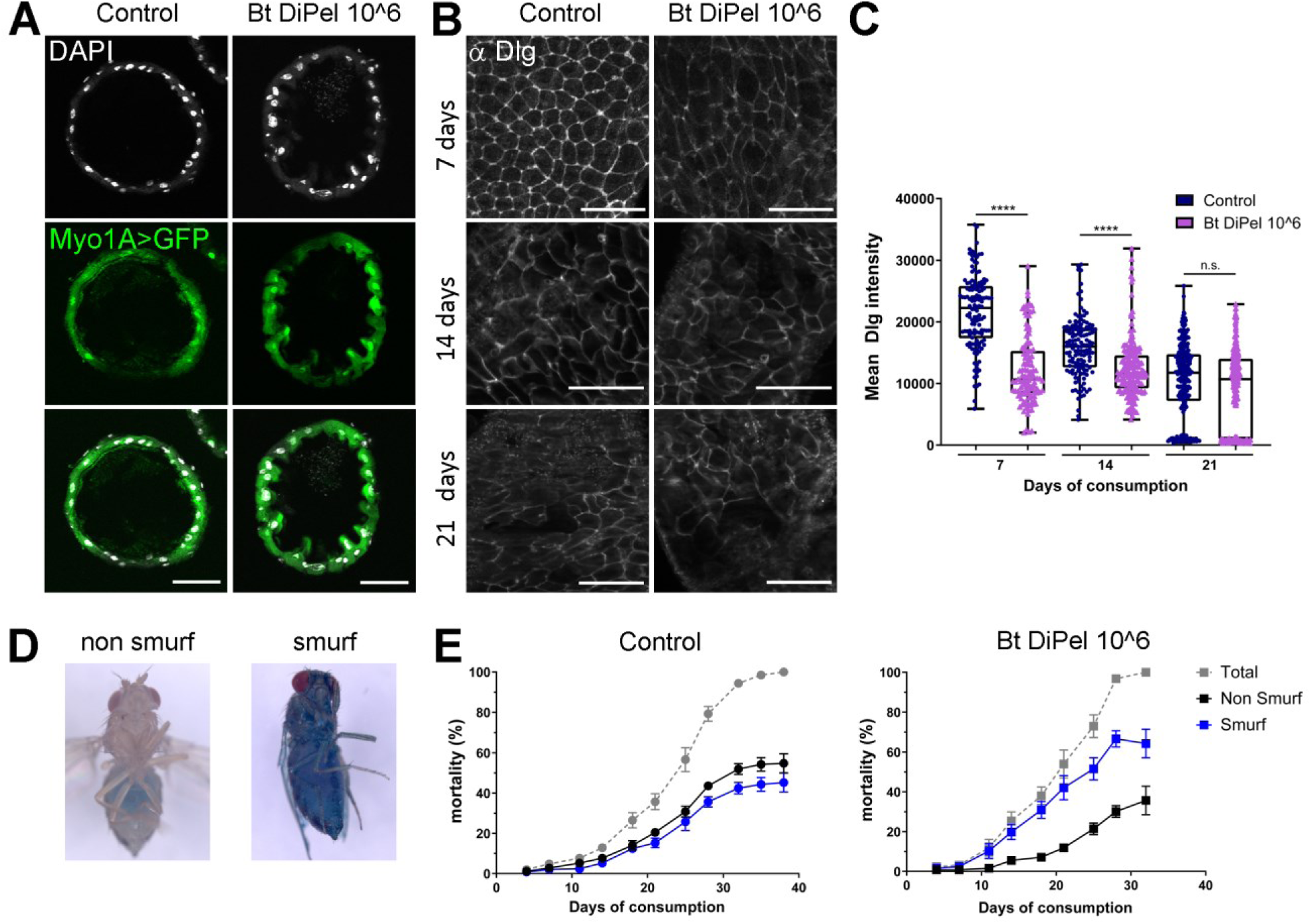
Enterocyte morphology, cell junctions and intestinal permeability after chronic *Bt* ingestion. **A)** Cross sections of intestines expressing GFP (green; Myo1A>GFP) in enterocytes at 14 days after DiPel exposure (10^6^ CFU/*Drosophila*/tube) compared to controls*. n=134 (Control 7d), n=112 (Bt 7d), n=136 (Control 14d), n=169 (Bt 14d), n=239 (Control 21d), n=233 (Bt 21d). Statistics were done with the Kruskal-Wallis test followed by the Dunn’s multiple comparisons test*. **B)** Immunostaining of Dlg septate junction protein over time (7, 14 and 21 days). **C)** Quantification of αDlg staining at enterocyte junctions**. D)** Illustration of the Smurf assay to assess intestinal permeability. Healthy non-smurf flies retain the blue dye in the intestinal lumen, while flies with loss of permeability have the dye diffused throughout the body (smurf). **E)** Quantification of the Smurf phenotype in flies treated with DiPel (10^6^) compared to control flies. *n=250 flies for the control and n=126 for DiPel*.

In the intestine, a decrease and cytoplasmic relocation of cell junction components increase gut permeability, and are tightly linked to aging and lethality in (55-57). We further investigated if morphological changes affect cell junctions and intestinal permeability upon chronic *Bt* ingestion. We first performed immunostaining for Discs large (Dlg), a protein localized to septate junctions, on posterior midguts at 7, 14 and 21 days. In control flies, Dlg staining nicely delineates the cell contours at day 7, and gradually diminishes during aging as previously reported (Fig 4B and C)(56). In contrast, we observed a strong loss of Dlg staining at septate junctions as early as day 7 in *Bt* conditions compared to the controls. At day 21, Dlg detection appears to be the same between *Bt* and control conditions. These results show that chronic *Bt* ingestion induces a premature alteration of midgut septate junctions that must occur naturally later with aging. As loss of cell junction components is detrimental to the epithelial barrier sealing, we assessed whether this was associated with a change in gut permeability by conducting Smurf assays (55). The assays revealed that the majority of flies treated with *Bt* present intestinal leakage unlike the control flies (Fig. 4D, E; Fig. S3). Taken together, the results show that chronic ingestion of *Bt* spores affects gut morphology, alters septate junctions and increases gut permeability, indicating the onset of early signs of midgut aging.

### Chronic *Bt* ingestion promotes progenitor dysplasia and raises ROS levels

Intestinal aging is also defined by inflammation-induced dysplasia, characterized by overproliferation of intestinal stem cells (ISCs) and mis-differentiation of progenitors, which naturally increase as flies age (58-60). To explore whether epithelial dysplasia occurs in the intestine of *Bt* treated flies, we analyzed both ISC proliferation rate and potential mis-differentiation of progenitors. First, we stained guts with the anti-phospho-histone 3 antibody (αPH3) to visualize the mitotic activity of the ISCs. In accordance with the literature, we observed an increase in ISC proliferation over time due to aging in control flies (Fig. 5A). A similar trend occurred in *Bt*-treated flies, however, the ISC proliferation rate appears to be significantly higher at each time point compared to control flies (Fig. 5A). We then quantified mis-differentiation, which is characterized by larger, rounded progenitors that form clusters of cells (58, 59). We used *esg-GAL4,UAS-GFP* flies (GFP expression in progenitors) and measured the proportion of GFP-positive progenitors forming clusters of more than 4 cells in control or *Bt*-challenged midguts (Fig. 5B). Control flies display increased mis-differentiation as time progresses, a trend also observed in *Bt*-treated flies. However, *Bt*-treated flies exhibit a statistically significant higher proportion of mis-differentiated clusters at days 14 and 21 (Fig. 5C). Altogether, these results indicate that chronic ingestion of *Bt* spores significantly exacerbates intestinal dysplasia, characterized by both proliferation and mis-differentiation of ISCs and progenitors, as early as 7 days after the onset of exposure.

**Figure 5.**
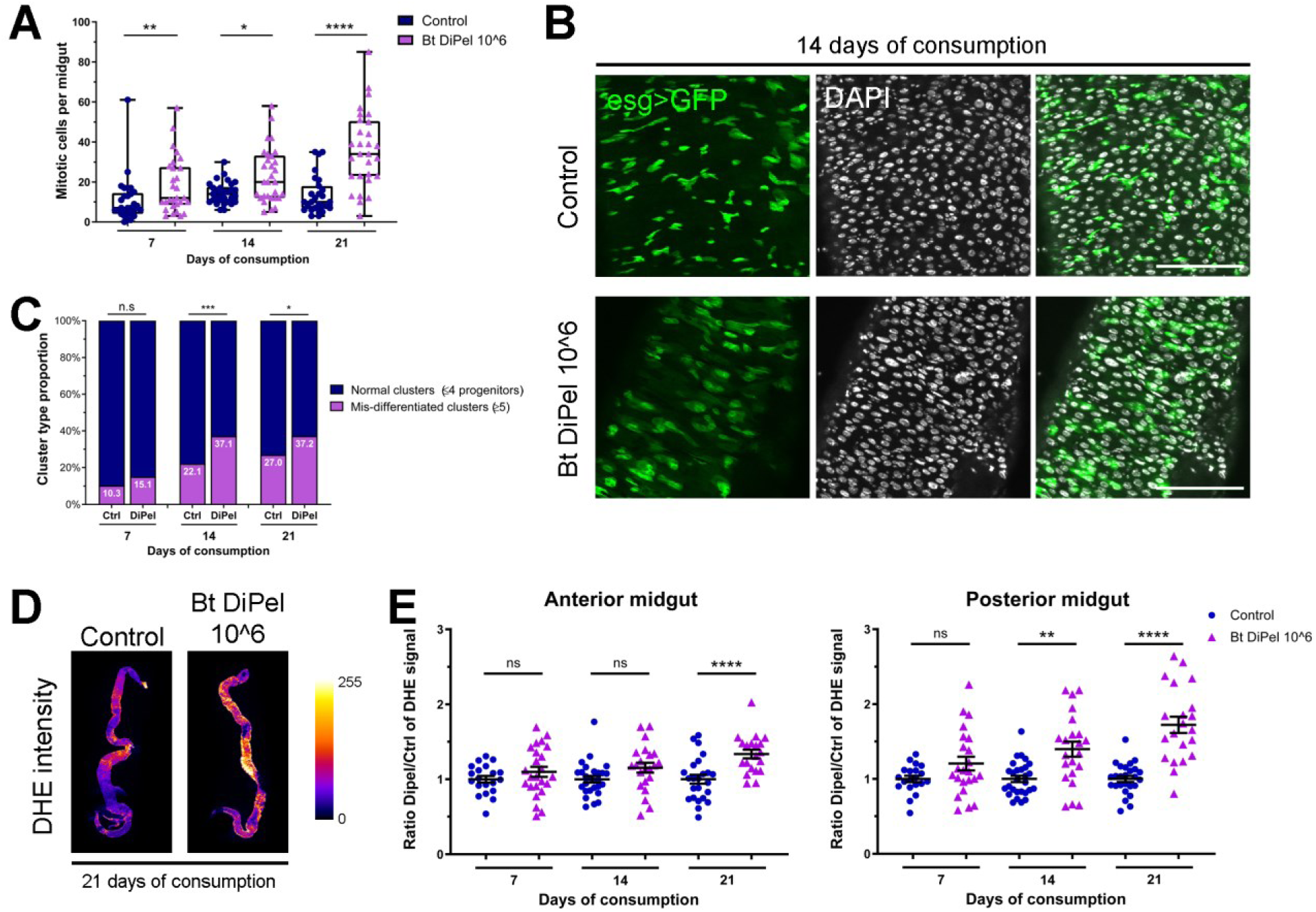
Dysplasia and oxidative stress after chronic *Bt* ingestion. **A)** Number of mitotic cells (αPH3 positive cells) per midgut in control and *Bt*-treated flies (10^6^ CFU/*Drosophila*/tube) at the different time points. *n=29 (Control 7d), n=29 (Bt 7d), n=30 (Control 14d), n=30 (Bt 14d), n=27 (Control 21d), n=29 (Bt 21d). Statistics were done with the Kruskal-Wallis test and the False Discovery Rate*. **B)** Images of *esgGAL4,UAS-GFP* control and *Bt*-treated intestines (at 14 days) expressing GFP in progenitor cells (GFP in green; DAPI in white). *n=175 (Control 7d), n=205 (Bt 7d), n=190 (Control 14d), n=202 (Bt 14d), n=204 (Control 21d), n=191 (Bt 21d). Statistics were done with the chi-square. test*. **C)** Quantification of progenitor clusters containing more than 4 cells (purple). **D)** Color gradient images (FIRE) showing representative intensities of DHE signal in control and *Bt*-treated intestines (at 21 days of consumption). **F)** Quantification of DHE signal over time after *Bt* treatment (control in blue and *Bt* DiPel at 10^6^ in purple) in the anterior and posterior parts of the midgut. *n=20 (Control 7d), n=24 (Bt 7d), n=27 (Control 14d), n=22 (Bt 14d), n=24 (Control 21d), n=20 (Bt 21d). Statistics were done with the Kruskal-Wallis test followed by the Dunn’s multiple comparisons test*.

Transcriptomic analysis pinpointed a consistent upregulation of genes involved in redox mechanisms. To support this result, we sought to detect the production of deleterious reactive oxygen species (ROS). We used the DHE probe to detect ROS as it strongly reacts with superoxide O2^−^, and to a lesser extent with other ROS (61, 62). Importantly, accumulation of high levels of mitochondrial ROS is a hallmark of aging (44, 63), and in *Drosophila*, old intestines exhibit elevated DHE-stained ROS levels (64). Compared to control conditions, *Bt*-treated flies show elevated DHE probe signal at day 21 in the anterior part and from day 14 in the posterior part (Fig. 5D and E). This result demonstrates that chronic *Bt* ingestion increases oxidative stress in the intestine. In addition, it is worth noting that the higher sensitivity of the posterior part to oxidative stress correlates with our previously published finding that *Bt* vegetative bacteria persist in this part of the midgut (31). Overall, our results show that chronic *Bt* ingestion enhances inflammation and oxidative stress, which accelerate gut aging and cause early lethality in *Drosophila*.

## DISCUSSION

*Bt* production of Cry entomopathogenic toxins is cleverly exploited as an insecticide to fight lepidopteran plant pests. The natural origin of the bacterial spores and the specificity of the Cry toxins have made it a favored choice over synthetic pesticides, contributing to more sustainable agricultural practices. As part of the *Bc* group, *Bt* shares many characteristics with the opportunistic *Bc sensu stricto*, which is responsible for foodborne outbreaks characterized by abdominal pains and diarrheal symptoms. Recent advances in genomics suggest that *Bt* and *Bc sensu stricto* belong to the same genomospecies (65). It is very difficult to distinguish them based on microbiological analyses solely and their genetic material has to be sequenced to make a distinction. The genomic similarity between the two species has raised concerns about whether the use of *Bt* as a biopesticide may contribute to food safety risks (19, 24-26, 66). In addition, the persistence of the spores and the presence of Cry toxins due to the high use of *Bt* products in agriculture increases the risk of their unintended effects on non-target organisms present in the environment (23).

In this work, we assessed the impact of chronic ingestion of *Bt* spores from commercial products using a non-target insect, *Drosophila melanogaster*. To do so, we mimicked plausible concentrations that may be encountered after spreading based on the recommendations within the European Union. We demonstrated that doses of 10^6^, and even 10^5^, CFU/*Drosophila*/tube reduce the longevity of flies and are detrimental to intestinal health. These doses are equivalent to two consecutive agricultural treatments and 1/5 of one treatment, respectively, indicating that low doses of *Bt* products could have unintended effects on *Drosophila*. Based on this information, the amount of *Bt* spores found in the environment could have long-term adverse effects on other non-target insects, which account for 85% of animal biodiversity. This suggests that the extensive use of *Bt*-based insecticides could destabilize ecosystems in the long term, and it is crucial to address this issue in the future to actively find appropriate solutions and reduce the risk associated with their ingestion.

The safety of *Bt* insecticides has been a topic of debate (18, 19, 66). Some studies suggest that the potential for *Bt* to act as a food safety hazard is minimal, while others emphasize the need for better differentiation between *Bt* and *Bc* in food safety evaluations. As *Bt* represents a major alternative solution to chemical pesticides and its use is increasing, its presence in the environment and on food items now requires more careful monitoring. A series of preventive and protective measures can be implemented, such as the dose and number of approved treatments, and more generally for consumers, the pre-harvest interval, the monitoring of *Bt* levels along the agri-food chain, and the systematic washing of fruits and vegetables.

## MATERIAL AND METHODS

### Reagents and resources

#### *Drosophila* strains

WT *Canton S* (RRID:BDSC_64349)

*w; myo1A-GAL4,UAS-GFP/CyO* (Y. Apidianakis)

*y w; esg-GAL4^NP5130^,UAS-GFP/CyO* (N. Tapon)

#### Bacterial strains

DiPel (DiPel® DF Jardin, Sumitomo Chemical Agro Europe)

Delfin (Delfin® Jardin, Certis USA)

*Bt kurstaki* ABTS-351 (18SBCL448, isolated from DiPel)

*Bt kurstaki* SA-11 (18SBCL487, isolated from Delfin)

*Bt kurstaki* SA-11Δcry (obtained from the strain SA-11 by plasmid curing (31))

*Bc Bactisubtil* (BGSC #6A8, (36))

*Bc* ATCC 14579 (collection NCBI, BGSC #6A5)

*B. amyloliquefaciens* H (collection NCBI, BGSC #10Al)

#### Antibodies

Anti-Disc large (DSHB #4F3-s)

Anti-Phospho-Histone 3 (Cell signaling 9701S)

#### Chemical and consumables

PBS-10X (Euromedex #ET330)

70% ethanol (VWR UN1170)

Triton X-100 (Sigma #T9284)

16% Formaldehyde solution Methanol-free (ThermoScientific #28900)

Fluoroshield^TM^ DAPI medium (SIGMA #F6057-20mL)

Erioglaucine disodium salt (Sigma #861146)

Elastase from porcine pancreas (SIGMA #E7885-5MG)

DHE (Dihydroethidium) probe (Invitrogen #D11347)

MicroElute Total RNA Kit (Omega #R6831-01)

#### Software and algorithms

FIJI/ImageJ (https://imagej.net/software/fiji); GraphPad (https://www.graphpad.com); Krita (https://krita.org/fr); R project (https://r-project.org)

### Experimental methods

#### *Drosophila* genetics

All *Drosophila* stocks were reared at 25°C on standard medium (0.8% agar, 2.5% sugar, 8% corn flour, 2% yeast) with a 12h light/12h dark cycle. For GFP expression, flies were crossed with WT to obtain a heterozygous F1 generation and to eliminate balancers for experiments. Flies of the desired genotype were collected within 24h after hatching and allowed to age for 7 days before starting any experiments.

#### Chronic *Bt* ingestion and *Drosophila* survival

10 flies (7 females and 3 males) were raised at 25°C on a vial containing fresh medium and waterman paper discs with 50µL of water (control condition) or *Bt* solution in order to give 1,25.10^6^ CFU/fly/vial of 4cm2 (*Bt* ingestion condition). We found that *Bt* spores placed on the nutrient medium in this way were starting to germinate from the fourth day (31). Flies were therefore transferred to new vials with fresh paper discs and solutions twice a week to avoid the presence of *Bt* vegetative bacteria coming from germinated spores. For all the experiments, only females were counted and dissected. At least three biological replicates were performed and males were added to replace dead females in order to maintain 10 flies per vial.

The recommendations for the spreading of *Bt* products in France are 1 to 1.5 kg/ha per application with repetitions ranging from 2 to 12 times ([E-Phy](https://ephy.anses.fr/); [EU Pesticides Database](https://ec.europa.eu/food/plant/pesticides/eu-pesticides-database/start/screen/active-substances)). This represents 1 to 1.5x10^13^ CFU/ha (10 000m^2^), equal to 4 to 6x10^5^ CFU/vial of 4 cm^2^, per application. The dose of 10^6^/fly/vial is therefore equivalent to 2 consecutive treatments.

#### Homemade spores

From isolated colonies on LB agar Petri dish, 4 x 5mL of *Bt* pre-culture was carried out. The pre-culture was used for sowing 4 x 500mL of PGSM medium (0.75% casamino acids, 0.34% KH2PO4, 0.435% K2HPO4, 0.75% glucose, 1.25mM CaCl2, 0.123% MgSO4, 0.002% MnSO4, 0.014% ZnSO4, 0.02% FeSO4) and allowed to grow and sporulate in an incubator shaker at 30°C, 180 rpm for 2 weeks. To eliminate vegetative cells, culture was heated 1 hour at 70°C and then centrifugated 15 min at 5000g. Each pellet were resuspended with a Dounce homogenizer in 250mL of 150 mM NaCl and placed for 30 min on roller agitation at room temperature. After centrifugation (15 min, 5000g), pellets were washed twice with sterile water (250mL). Final pellets were resuspended in 30mL of sterile water, dispatched in 1mL weighed tubes and lyophilized for 24-48h. The spore mass was determined by the difference between the full and the empty tubes weights.

#### Quantification of intestinal bacterial load

Female *Drosophila* were washed for 20s in 70% ethanol prior to gut dissection in PBS 1X. Only the proventriculus and midgut were retained and then ground in 400μL LB using a micro-pilon. The ground intestines were then divided into two tubes, one to quantify the total bacterial load in the intestine and another to quantify intestinal spores. For spore quantification, samples were heated at 75°C for 25 min to eliminate vegetative bacteria. Each tube was then serially diluted (10^-1^ to 10^-3^) in LB and incubated at 30°C on LB agar plates. Colonies were counted the following day. The difference between the total bacterial load of the unheated sample and the amount of spores in the heated sample was used to determine the amount of vegetative gut bacteria. These experiments were carried out on days 7, 14 and 21 of intoxication.

#### Smurf test: monitoring of intestinal permeability

Smurf assays were performed as described (55). Briefly, *Bt* exposure was performed as described above, except that a water solution containing a food dye (Erioglaucine disodium salt 5%) was used to resuspend *Btk* spores (ABT-351 or DiPel). Mortality of Smurf females and non-Smurf females was counted every 3-4 days. As in the longevity test, dead females were replaced by males to keep the number of flies per tube constant.

#### Immunochemistry and quantification

After dissection in PBS 1X, intestines were fixed in formaldehyde for 1h and then blocked in a solution of BSA% + PBS-Tween 0.2%. Anti-Dlg and anti-Phospho-Histone 3 (PH3) primary antibodies were used at 1:200 dilution and 1:500 dilution, respectively. The intestines were then washed, labeled with the secondary antibody and mounted with Fluoroshield^TM^ medium containing DAPI. Images are acquired using the AxioImager Z1 with Apotome 2 or LSM880 microscope (Zeiss).

For Dlg quantification, experiments were performed in triplicate or quadruplicate, with an average of 5 intestines and about 40-45 tight junction measurements per replicate. Maximum intensity projections of the images were done. The mean intensity of the control was calculated for each replicate, and the ratio of this mean to the sample value was calculated. For the PH3 quantification, cells stained with the anti-Phospho-Histone 3 antibody were counted in the whole midgut. For the evaluation of progenitor mis-differentiation, GFP-labeled clusters (esg>GFP) were manually circled with FIJI and the number of associated nuclei was counted.

#### DHE staining and quantification

DHE staining was performed as previously reported (61). Intestines were dissected in 1X Schneider’s *Drosophila* medium. Images were acquired with the axio Imager Z1 equipped with the apotome 2 module and recorded in sum intensity projection. Experiments were performed in triplicate, with approximately twenty intestines analyzed at a time. For each intestine, the mean intensity was calculated separately for the anterior and posterior sections. For each intestinal section, the mean intensity value of the control was determined and a ratio was calculated between this mean value and that of the sample.

#### RNA sequencing

Midguts of mated female flies were dissected in PBS-1X and homogenized with a TissueLyser LT (Qiagen) using 5 mm beads at 50 hz for 50 seconds. Total RNA was extracted using the MicroElute Total RNA Kit (Omega) according to the manufacturer’s instructions. Three independent replicates were carried out at each time point (7d, 14d, 21d). RNA sequencing was performed by Genewiz (Azenta Life Sciences) using Illumina NovaSeq, PE 2x150 technology (version 2.0.0) in a polyA pair-end manner.

Raw data were mapped to the *Drosophila melanogaster* BDGP6 reference genome (dmel_r6.32) available at ENSEMBL, and are currently being deposited on the European Nucleotide Archive ([ENA](https://www.ebi.ac.uk/ena/browser/home)). Gene hit counts were used for downstream differential expression analysis using “EBSEQ”. The Volcano plot was generated using the EnhancedVolcano in R library. Gene Ontology analysis was performed on the differentially expressed genes of the three time points regrouped using the *enrichGO* function from the clusterProfiler R package, and plots were produced using the ggplot2 package.

#### Statistics

For all experiments, flies were randomly chosen for experimental analysis. All experiments were independently repeated at least 3 times. Statistical tests were performed with R software or GraphPad. The total number of samples (“*n*”) and the tests used are indicated in the legend of each figure.

## ACKNOWLEDGEMENTS

We wish to thank Nicolas Tapon, Yiorgos Apidianakis, the Bloomington Drosophila Stock Center (BDSC; NIH P40OD018537) and the Developmental Studies Hybridoma Bank (DSHB, created by the NICHD of the NIH and maintained at the University of Iowa, Department of Biology, Iowa City, IA 52242) for fly stocks and reagents. We are very grateful to the Alain Robichon Foundation for the financial support of the RNA-seq. We acknowledge Olivier Pierre from the imagery platform of our “Institut Sophia Agrobiotech” and Julie Cazareth from the flow cytometry and cell sorting platform of the Institute of Molecular and Cellular Pharmacology (Sophia Antipolis) for their assistance. We are grateful to the bioinformatics and genomics platform, BIG Sophia Antipolis (ISC plantBIOs, https://doi.org/10.15454/qyey-ar89), for computing and storage resources. We thank all the members of our team “*Bacillus*, Environment and Health” for insightful discussions. A.J. was supported by the “Ligue contre le cancer” (doctoral grant 2018) and the “Fondation ARC pour la recherche sur le cancer” (ARCDOC42021020003051). J.F. is supported by a PhD grant from INRAE (IB2024 n° 9695) and the action under Ecophyto II+ plan (BaThuThese). R.R., A.G. and C.R. are supported by the “Centre national de la recherche scientifique” (CNRS). This work was funded by the “Fondation ARC pour la recherche sur le cancer” (20171206145), by the PNR-EST ANSES & ECOPHYTO II (2017), by the “Institut Olga Triballat” (AAP2021; institut-olgatriballat.org), and by the ANR-564 22-CE35-0006-01 (BaDAss). We were also funded by the action under Ecophyto II+ plan, led by the Ministries of Agriculture, Ecology, Health and Research, with financial support of the French Office for Biodiversity (BaThuGut). In addition, we received support from the French government, through the UCAJEDI Investments in the Future project managed by the National Research Agency (ANR) with the reference number ANR-15-IDEX-01 (AAP : AO1 Fonctionnement 2022, Space, Environment, Risk and Resilience Academy 3 of Université Côte d’Azur).

---

**Supplementary Figure S1.**
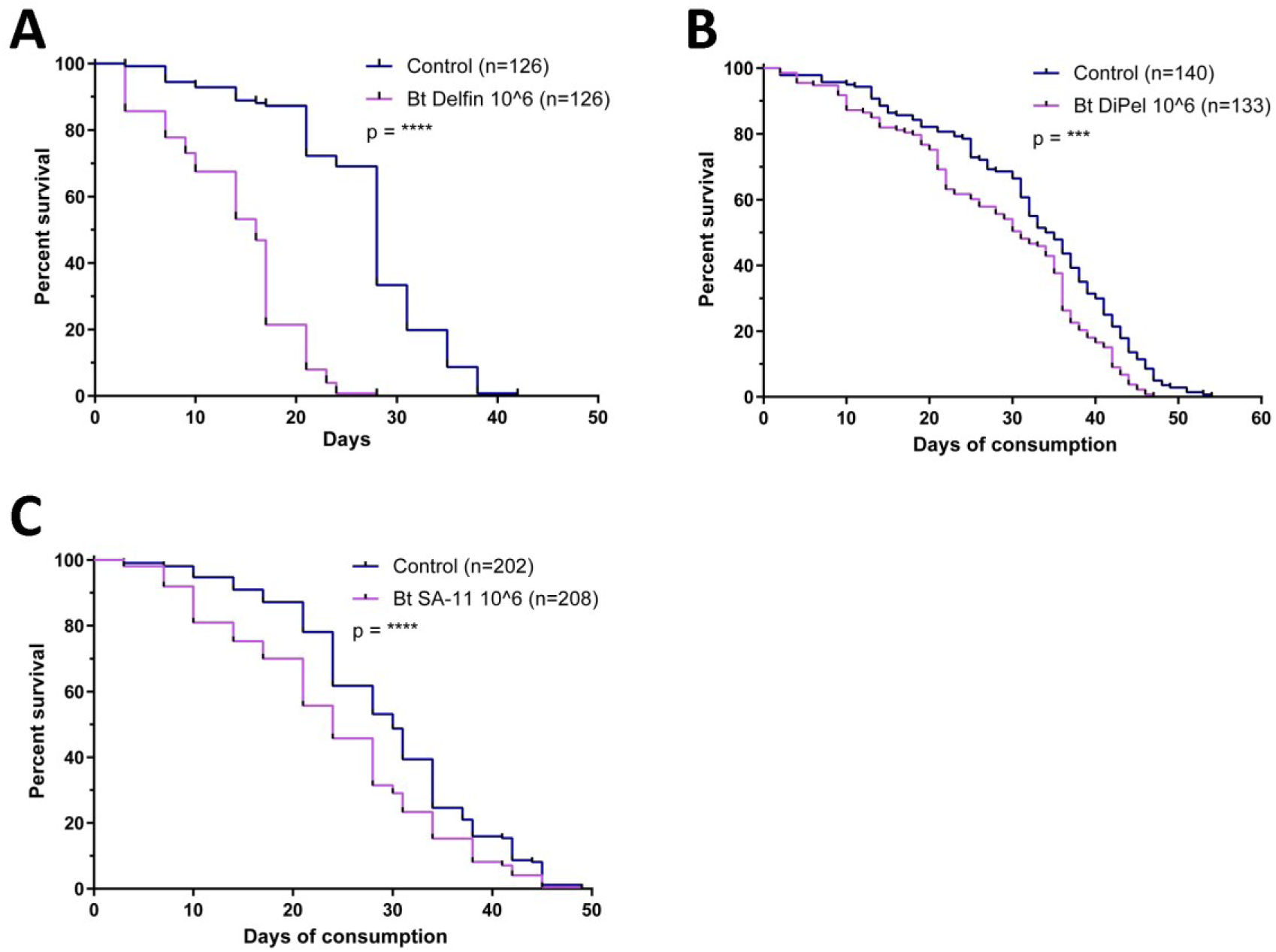
*Drosophila* survival after chronic ingestion of *Bt* spores. **A**) Survival after ingestion of the Delfin commercial product. **B)** Survival assay when vials were changed every day. **C)** Survival after ingestion of homemade *Bt* SA-11 spores. *Doses and n are indicated in each panel and results shown as Kaplan-Meier curves with Log-Rank Mantel-Cox test*.

**Supplementary Figure S2.**
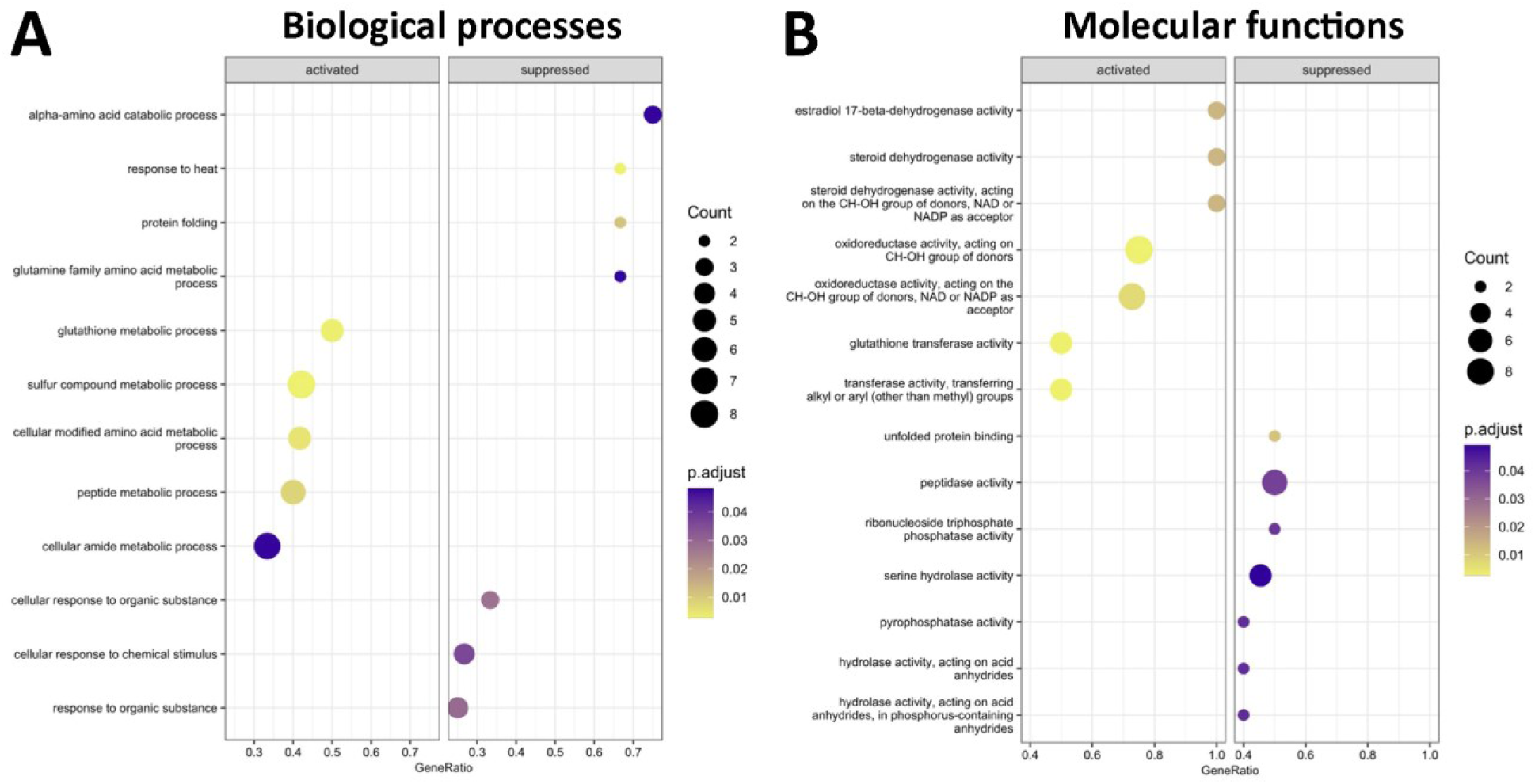
GO terms found in differentially expressed genes. Representative Gene Ontology terms enriched in upregulated (activated) and downregulated (suppressed) genes in the intestines of *Bt*-treated flies compared to control flies (all time points regrouped). **A)** Biological processes and **B)** Molecular functions.

**Supplementary Figure S3.**
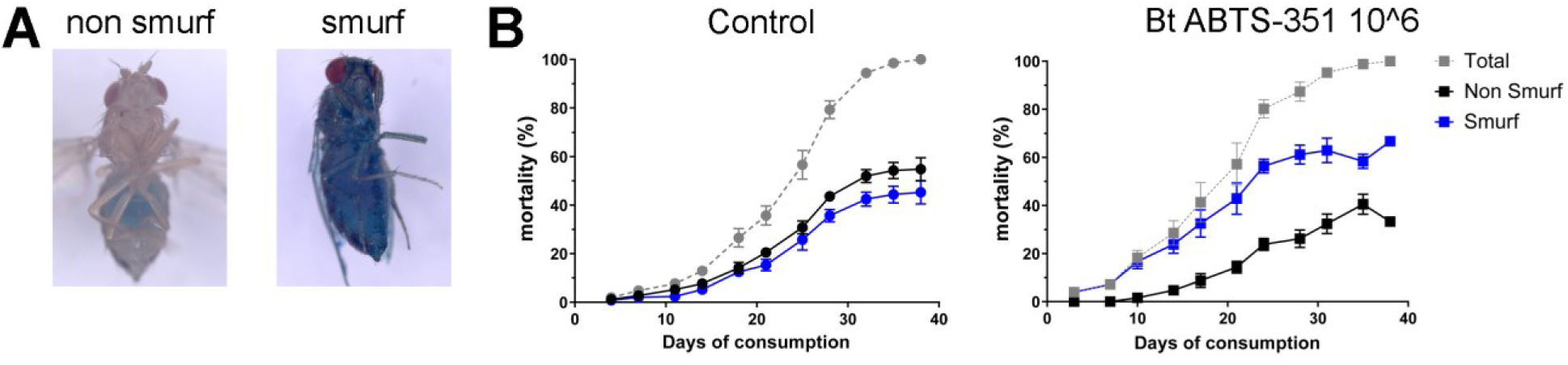
Smurf assays. **A)** Illustration of the Smurf assay to assess intestinal permeability. Healthy non-smurf flies retain the blue dye in the intestinal lumen, while flies with loss of permeability have the dye diffused throughout the body (smurf). **B)** Quantification of the Smurf phenotype in flies treated with the laboratory produced DiPel-derived ABTS-351 spores (10^6^) (without additives) compared to control flies. *n=250 flies for the control and n=126 for DiPel*.

